# A Hot-Swappable Genetic Switch: Building an inducible and trackable functional assay for the essential gene MEDIATOR 21

**DOI:** 10.1101/2024.12.16.628800

**Authors:** Isabella J. Watson, Cassandra Maranas, Jennifer L Nemhauser, Alexander R Leydon

## Abstract

Essential genes, estimated at approximately 20% of the *Arabidopsis* genome, are broadly expressed and required for reproductive success. They are difficult to study, as interfering with their function leads to premature death. Transcription is one of the essential functions of life, and the multi-protein Mediator complex coordinates the regulation of gene expression at nearly every eukaryotic promoter. In this study, we focused on a core Mediator component called MEDIATOR21 (MED21), which is required for activation of transcription. Our previous work has also shown a role for MED21 in repression of gene expression through its interaction with a corepressor protein. Here, we sought to differentiate the role MED21 plays in activation versus repression using the model plant *Arabidopsis*. As mutations in MED21 lead to embryo lethal phenotypes, we constructed a set of synthetic switches using PhiC31 serine integrases to create an “on-to-off” inducible loss of function MED21 in a non-essential tissue. Our technology, which we call Integrase Erasers, made it possible for *med21* mutant plants to survive into adulthood by ablating protein expression selectively in lateral root primordia, allowing quantification and characterization of *med21* mutant phenotypes in a post-embryonic context. In addition, we engineered chemical induction of the Integrase Eraser to ablate MED21 expression in whole seedlings at a user-specified timepoint. Finally, we extended this technology to build a hot swappable Integrase Isoform Switch where expression of the integrase toggled cells from expressing wildtype MED21 to expressing MED21 sequence variants. Our analysis of the entire set of new integrase-based tools demonstrates that this is a highly efficient and robust approach to the study of essential genes.

## Introduction

Essential genes carry out functions required for survival. They are also likely to be broadly expressed, show up as hubs in protein networks, and be components of stable complexes with other biomolecules^1–3^. Examples like general transcription factors (GTFs) and the Mediator complex play significant roles in regulating transcription at most genes via interactions with gene-specific DNA-binding transcription factors^4–7^. The difficulty in assigning function to Mediator subunits is well-documented in plants^4,5,8,9^.

Previously, we reported that activity of the plant corepressor TOPLESS (TPL) depends on interaction with the Mediator complex through direct contact with the middle domain subunit MEDIATOR21 (MED21)^10,11^. One consequence of the embryo lethal phenotype of *med21* loss of function mutants^12^ has been difficulty differentiating its mechanism of action during gene activation versus gene repression.

There are several approaches that have been employed to study essential genes in yeast, such as temperature sensitive alleles^13^, chemically induced nuclear depletion (anchor away)^14^, and auxin induced degradation (AID)^15^. Adaptations of these approaches have been employed in plants^16–18^, yet the majority of essential genes have been characterized by the embryo lethal phenotypes found in T-DNA insertion mutants^19^. Recent advances in the application and optimization of serine integrase activity in *Arabidopsis thaliana* provides an alternative means to efficiently, precisely and irreversibly alter relatively large genomic regions.

Serine integrases, hereafter referred to as integrases, induce recombination events at asymmetric DNA sequences that are specific to each integrase. Two integrases, Bxb1 and PhiC31, were recently optimized to build a modular toolkit for *Arabidopsis*^20^. In that work, the integrase target site contained a ubiquitous promoter flanked by integrase sites, such that expression of the cognate integrase switched expression from one fluorescent protein to another. Further work using more complicated target design and cell-type specific developmental promoters allowed order-of-expression cell lineage tracing^21^.

Here, we extended the work on initial integrase prototypes^20^, to study the essential gene *MEDIATOR21* (*MED21*). We constructed an “on-to-off” inducible loss of function integrase target, which we term an Integrase Eraser. This target switches expression from MED21 to a fluorescent reporter thereby permanently labeling cells with altered genomes. We tested target behavior using two different PhiC31 expression schemes *in planta*: 1) an integrase driver that induces loss of function specifically in lateral root primordia, and 2) an inducible integrase driver that enables synchronized switching in all cells at a given timepoint. By expressing these constructs in a *med21* mutant plant in combination with the MED21 target, both approaches successfully circumvent the embryonic lethal phenotype. In addition, we engineered an “isoform switch” integrase target, where MED21 is expressed as a P2A fusion with a fluorescent protein, differentially marking cells expressing wild-type or mutant isoform. This new collection of tools, and the ready adaptation of its components to most contexts of potential interest, unlocks the study of essential genes in post-embryonic plants, while specifically advancing our understanding of the Mediator complex in coordinating transcriptional regulation.

## Results and Discussion

To better understand the function of *MED21* in *Arabidopsis*, we built a synthetic circuit that would allow for selective loss of *MED21* function in lateral roots, a non-essential organ. This Integrase Eraser technology is composed of two parts: an “on-to-off” inducible loss of function integrase target and a driver line that expressed the PhiC31 integrase from a cell-type-specific promoter, pGATA23. To build our target, we adapted a golden gate assembly approach from Guiziou et al., 2023^20^ that yielded a final construct with a constitutive promoter (pUBQ10) flanked by integrase sites, positioned between a wildtype copy of *MED21* fused to an HA epitope tag and a reporter gene (mScarlet) (Fig. 1A). We also cloned a version of the target lacking the HA tag to control for effects of tag placement on the MED21 protein. Both constructs were transformed into a heterozygous T-DNA loss of function *med21-1* mutant. The target constitutively complements the loss of MED21 until the integrase is active, so it is transformed into mutant plants before introduction of the driver line (Figure 1B). All cells where the integrase mediates an inversion in the target (a “switch”) are marked by expression of mScarlet (Figure 1B, lower arrows). Therefore, the presence of mScarlet indicates the loss of wild-type *MED21* activity (Fig. 1B).

**Figure 1.**
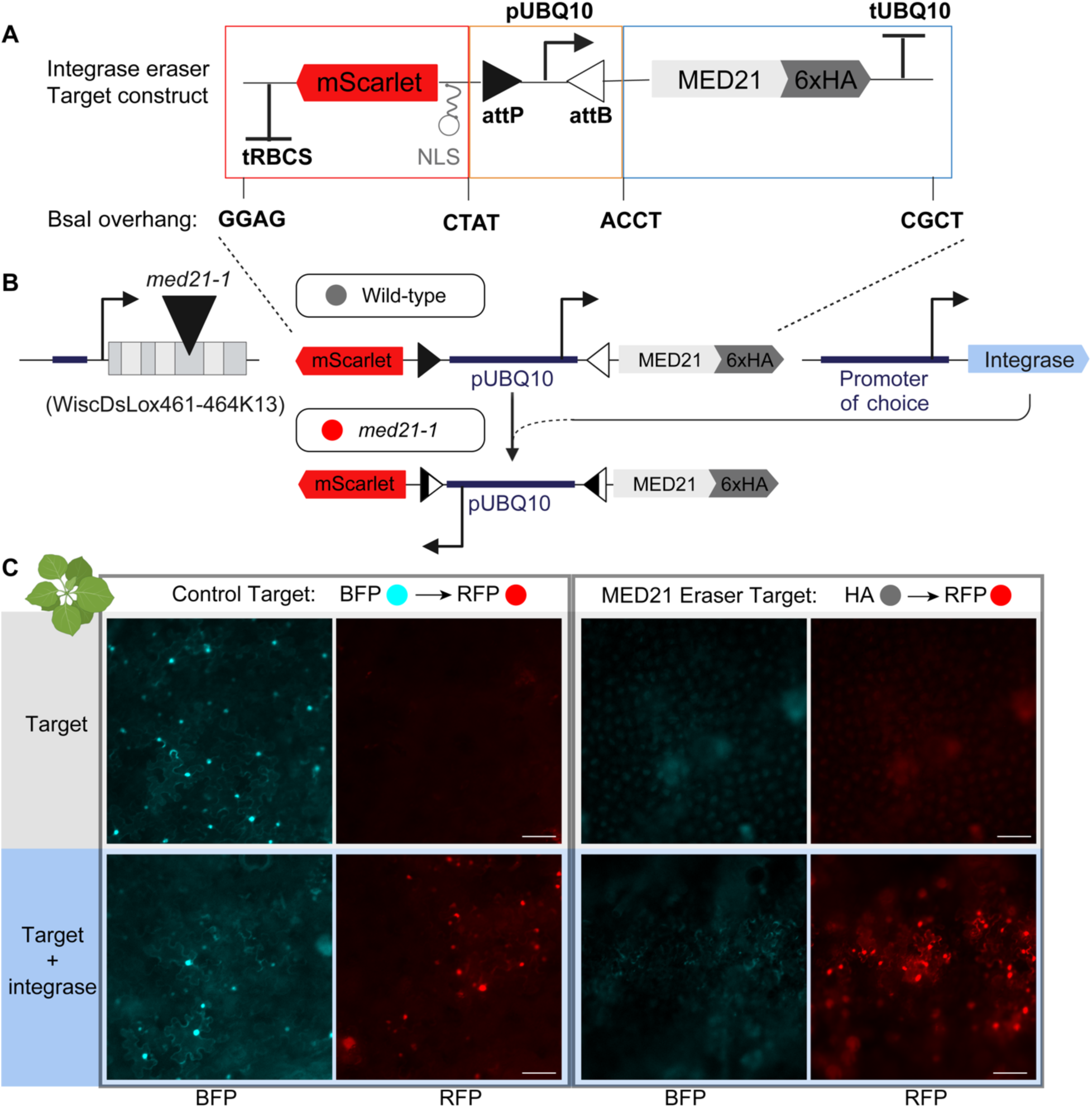
Engineering and rapid prototyping of the integrase eraser approach. **A.** Cloning strategy based on (Guiziou et al., 2023^20^) for the target integrase target with BsaI adapters highlighted for use in golden gate cloning strategies. **B.** Design of the integrase target. The target is composed of two PhiC31 integrase sites (triangles) surrounding a constitutive promoter (pUBQ10), the rescue coding sequence for MED21:6xHA and the fluorescent reporter (mScarlet). In absence of integrase, MED21:6xHA is expressed. In presence of integrase, the integrase mediates inversion of the DNA between the integrase sites, inverting the promoter, and leading to mScarlet expression. The expression of the integrase is mediated by the promoter selected. **C.** Rapid prototyping of the MED21 Eraser target in transient transfections of *Nicotiana benthamiana* at 2 days after injection. On the left side, a control target that switches from mTurquoise to mScarlet alone (top) and with a *p35S:PhiC31* construct (bottom). On the right side, the MED21 target alone (top), and MED21:HA target with a *p35S:PhiC31* construct (bottom). The BFP channel for the MED21:HA is shown to demonstrate the level of background expected for the BFP channel in the negative control. Microscopy images were taken on a 20x objective to allow a wide view of switching efficiency, and the 50µm scale bar in the RFP channels applies to all paired BFP images.

We used rapid prototyping in *Nicotiana benthamiana* to test the functionality of the MED21 Integrase Eraser target. As a control, we used a previously characterized target that switches from mTurquoise to mScarlet in the presence of the PhiC31 integrase expressed from the strong viral 35S promoter (Figure 1C). Without the addition of PhiC31 integrase, no mScarlet was detected for either control or MED21 Eraser targets, indicating no stochastic switching occurred. In contrast, in the presence of PhiC31, we observed the presence of cells that expressed nuclear mScarlet. These results confirmed that the MED21 Integrase Eraser was functional and specific.

Next, we generated stable transgenic *Arabidopsis* plants expressing the *MED21* Integrase Eraser constructs. We first focused on erasing *MED21* activity during lateral root initiation, a well-characterized example of de novo organogenesis (Banda et., al 2019). Lateral roots are an attractive candidate for this Integrase Eraser technology, as they are not required for plants to survive in controlled environments like growth rooms or greenhouses. Thus, the impact of losing essential gene function can be studied while maintaining largely wild-type levels of plant health and reproductive fitness. Lateral roots develop from a small subset of the xylem pole pericycle (XPP) termed founder cells.

*GATA TRANSCRIPTION FACTOR 23 (GATA23)* is strongly and largely specifically expressed in these founder cells, setting into motion the series of asymmetric cell divisions that form the new root^22^. Previous use of a *pGATA23* integrase driver produced many lines with switching only in cells in the lateral root, that is, cells from a lineage involving *GATA23* expression^20^. Our use of this same promoter in the *MED21* Integrase Eraser should “erase” *MED21* in all of the cells that compose the lateral roots (Fig 2A, red lateral roots). Importantly, in all MED21 eraser lines, we observed no detectable expression of the upstream RFP gene by microscopy (0/43 MED21:HA target, 0/49 MED21 target) in plants that do not carry an integrase expression construct, demonstrating that pUBQ10 is unlikely to act as a strong bidirectional promoter, consistent with our other implementations in *Arabidopsis*^20,21^.

**Figure 2.**
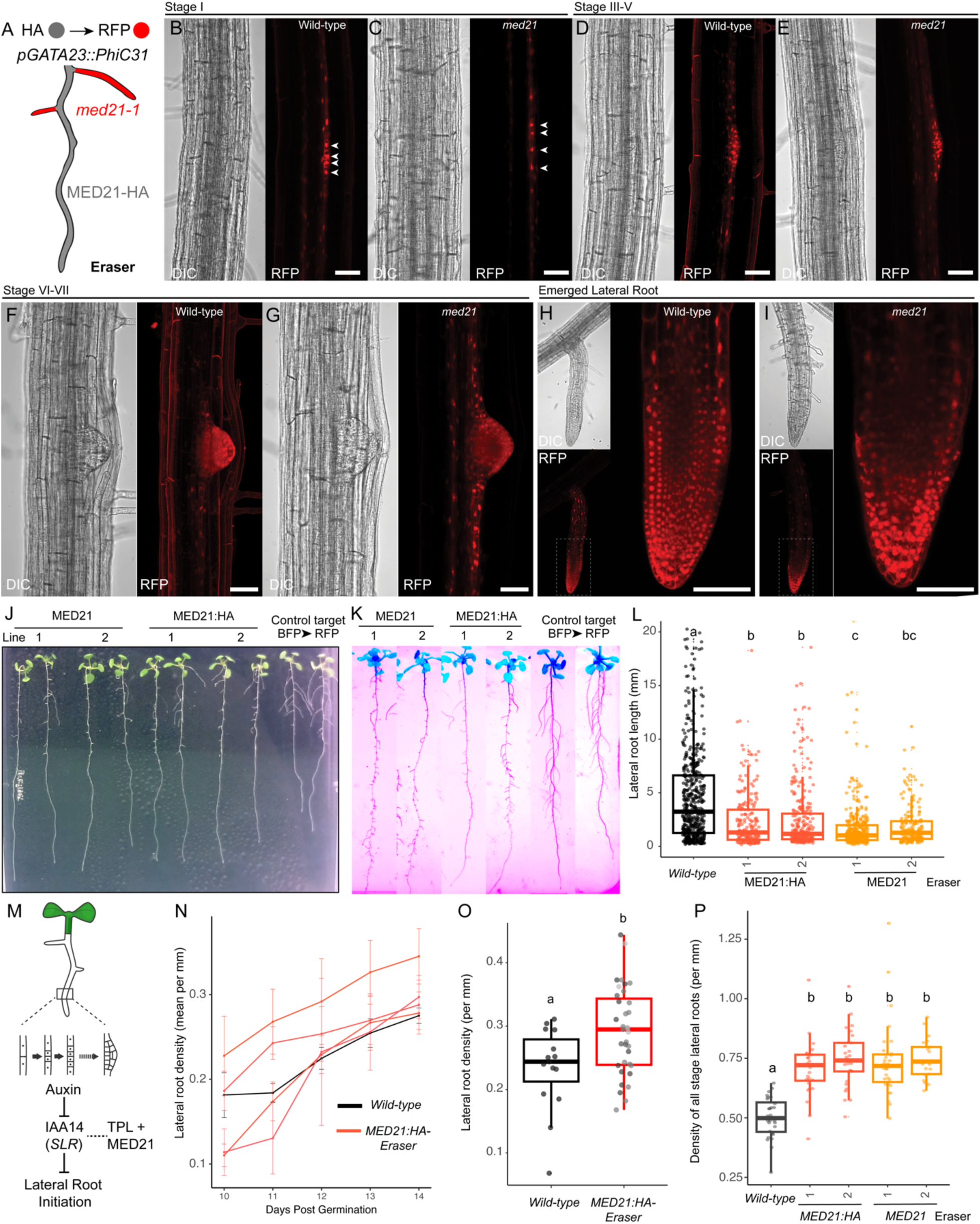
Cell type specific integrase eraser implemented in lateral root primordium results in *MED21* loss of function and increases in lateral root initiation. **A.** Schematic of the predicted behavior for the MED21 eraser approach. Lateral root primordium cells (red) will lose expression of the rescue construct, rendering them an effective knockout for MED21. **B-J.** Confocal microscopy analysis of wild-type control switch (from Fig. 1C) and MED21 eraser lateral root initiation. All scale bars are 50µm. **J.** Still image from Supplemental Movie 1 of the root growth phenotype of MED21 eraser lines compared to wild type at day 11. **K.** Whole seedling epi-fluorescent image of MED21 eraser lines at day 14. Images have been color inverted to allow visualization of the short root phenotype. Natural autofluorescence was captured for whole seedlings on plates allowing all root tissue to be observed. **L.** Lateral root lengths quantified at day 14 from individual selected T2 lines for each eraser type.

After introducing the *pGATA23::PhiC31* driver into *med21-1* plants complemented by the MED21 eraser target (Fig. 1B), we identified lines with mScarlet expression restricted to the lateral roots (Figure 2B; 5 independent MED21:6xHA eraser lines and 9 independent untagged MED21 eraser lines), and compared these to the control switch (Fig. 1C) through confocal microscopy. All selected lines demonstrated essentially wild-type pre-emergence lateral root development (Figure 2B-G; stages I-VII). However, the loss of MED21 did lead to mild disruptions of architecture in post-emergence roots. As early as 10 days post germination (dpg), we observed that MED21 eraser lateral roots had larger root cap zones and abnormally shaped root hairs (Figure 2H-I), as well as a reduced length compared with our wild type controls (Figure 2J). The latter phenotype was even more striking at 14 dpg (Figure 2K,L). Dynamic tracking of lateral root development over 16 days (Supplemental Video1) highlights the impact of loss of MED21 activity. We observed switching primarily from the early stages of the lateral root initiation, however in future implementations the pGATA32 promoter could be tuned to reduce integrase activity to improve specificity through the inclusion of protein degrons to the integrase, or RNA destabilization tags (DSTs) to its transcript^20^.

Previous work has shown that the corepressor TPL interacts with the Mediator complex directly through contacts with MED21^10^ (Figure 2N). The interaction between the Mediator complex and TPL is critical in maintaining repression of transcription in yeast and plants^10,11^. One of the best-studied TPL-regulated pathways is auxin response^23^. In low auxin environments, the TPX family (TPL/TPRs) inhibits the activity of Auxin Response Factor (ARF) transcriptional activators through their mutual binding of Aux/IAA proteins (AUXIN/INDOLE-3-ACETIC ACID). When auxin levels rise, Aux/IAA proteins are degraded, disrupting the connection between ARF and TPX proteins, thereby activating expression of ARF target genes^24^. Following this logic, loss of MED21 should reduce TPL activity and phenocopy the addition of auxin addition. In lateral roots, loss of MED21 should therefore promote initiation, an auxin-regulated process (Figure 2M). To test our model, we performed a time course growth analysis of MED21:HA eraser lines to determine whether lateral roots were initiated more frequently than controls (Lateral root density, Figure 2N). We observed that over time, the MED21:HA eraser lines did indeed have a higher lateral root density than wild-type controls (Figure 2N,O). When we included pre-emergence lateral roots, this trend was even more clear (Figure 2P).

Experiment was performed in triplicate for each genotype, with 6 seedlings per plate (n=18 per genotype, at least 180 lateral roots per genotype). **M.** Auxin-induced degradation of the IAA14 is strictly required for initiation of lateral root development. IAA14 recruits TPL which in turn inhibits the mediator complex through interaction with MED21. Cartoon is modified from^10^. **N.** Lateral root density over time for MED21:HA eraser lines. Data is presented as means +/- standard error. Experiment was performed in triplicate for each genotype, with 6 seedlings per plate (n=18 per genotype). **O.** Lateral root density at day 14 of growth for MED21:HA eraser lines. Each gray color represents an individual lateral root density from 4-6 independent lines and is representative of the day 14 data from N. **P.** Lateral roots of all stages (emerged and non-emerged) were quantified in homozygous mutant lines at day 14. For **L,O,P.** Letters indicate significant difference (ANOVA and Tukey HSD multiple comparison test; p<0.001).

To enable loss of MED21 activity throughout the plant at a specific timepoint, we engineered a chemically-inducible *MED21* Eraser (iEraser) (Figure 3A). To do this, we replaced *pGATA23* with the well-characterized β-estradiol system, as it is small, fast-acting in nearly every plant cell after treatment, and is fully orthogonal to known plant hormones^20,25^. We used our *MED21:HA* target for these experiments, as it functioned similarly to untagged *MED21* (Figure 2) and makes it possible to assay protein levels. Complemented *med21-1*, MED21:HA Eraser lines were transformed with *p𝛃-estradiol::PhiC31*. Progeny from selected transformants (T2 plants) were transplanted onto estradiol plates for analysis. By 48 hours, we observed strong and specific induction of MED21:HA switching only in seedlings exposed to β-estradiol (Figure 3B-C). We defined a seedling as having switched when nuclear RFP was detected in the root tip region by microscopy, and consider a line specific when all roots show at least some cell switching (see methods). Previous integrase induction by β-estradiol was detectable around 48 hours, with more than 50% of seedlings fully switched by 72 hours^20^. In our experiments, we detected 100% switching by 48 hours. The efficiency of switching in both tagged and untagged MED21 iEraser lines was similar, with little non-specific switching found (Figure S1A). The rate of uninduced lines demonstrating leaky switching is consistent with our own observations that the XVE system has a low but detectable transcriptional leakiness and highlights the importance of thorough screening of multiple independent lines. MED21:HA iEraser seedlings demonstrated increased root hair development (Figure 3B-C) and growth reductions following estradiol treatments compared to controls lacking β-estradiol (Figure 3D).

**Figure 3.**
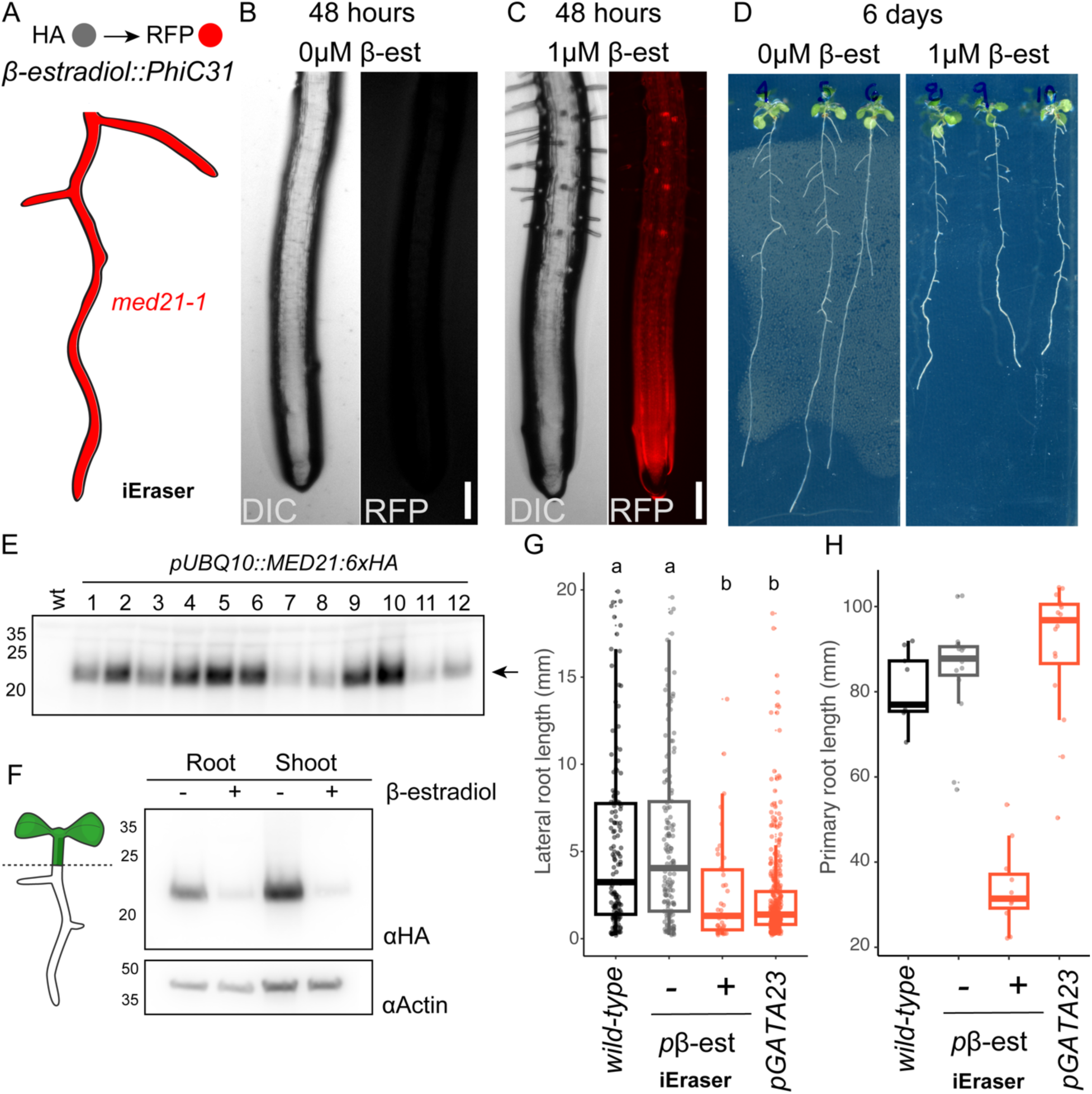
A chemically inducible MED21 eraser. **A.** Schematic of the predicted behavior for the MED21 iEraser. The estradiol inducible integrase construct is composed of p35S:XVE (transcriptional activator composed of a DNA-binding domain of LexA, the transcription activation domain of VP16, and the regulatory region of the human estrogen receptor;) and pLexA-minimal 35S driving expression of PhiC31. **B-C.** Characterization of the iEraser by fluorescence microscopy shows induction as early as 48 h after treatment. Scale bar represents 50μm. (**C.**). **D.** Growth phenotypes were visible in estradiol induced switched plants grown on 1µM 𝛃-estradiol induction plates for 6 days. Seedlings were first grown for 6 days on LS media lacking 𝛃-estradiol before transplanting to induction media (12 days total growth). **E.** Protein expression analysis by western blot for twelve independent iEraser lines. **F.** Protein expression analysis by western blot for roots and shoots isolated from plants grown with or without 𝛃-estradiol. Seedlings were grown on LS media for 6 days, and then transplanted to either control or induction media for another 6 days of growth. Experiment was performed in triplicate for each genotype. **G-H.** Lengths of lateral **(G)** and primary **(H)** roots were quantified at 6 days from the indicated Integrase Eraser type. Seedlings were grown as in **F**. Letters indicate significant difference (ANOVA and Tukey HSD multiple comparison test; p<0.001).

We characterized the expression of *MED21* in twelve independent iEraser lines, using the HA epitope tag. All lines had detectable protein expression (Figure 3E), so we focused further characterization of the MED21 iEraser on the progeny of one of these lines. Loss of MED21:HA could be detected in root and shoot tissues isolated from iEraser seedlings exposed to β-estradiol for 6 days, although a low level of protein was still visible on western blots (Figure 3F). It is possible that the UBQ10 promoter drives high MED21:HA protein levels and therefore longer MED21:HA turnover times, and this could be tuned down by the addition of RNA destabilization tags to achieve lower abundances^20^. This raises the possibility that β-estradiol may not be reaching a subset of cells, as has been seen for certain cell types like meristematic tissues^26,27^. To test this hypothesis, we took advantage of the irreversible nature of the integrase-mediated switch. We reasoned that if all cells were switched in a parent plant, then the cells of the subsequent gametophyte and germ cells would also be switched, leading to a fully-switched seedling in the next generation. Instead, we observed no switching in the progeny of seedlings exposed to β-estradiol (Figure S1B), indicating that at least a subset of meristematic cells must escape induction of PhiC31.

Significant growth reduction in both primary and lateral roots could be measured in MED21 iEraser seedlings treated with β-estradiol (Fig 3D). This phenotype is consistent with the predicted impact of loss of this essential gene. As expected, we found that *pβ-estradiol::PhiC31* phenocopied the reductions in lateral root length (Figure 3G) observed in *pGATA23::PhiC31*. Consistent with the difference in expected expression patterns of each promoter, *pGATA23* Eraser lines had similar primary root length to wild type, while primary root length in iEraser plants was sharply reduced (Figure 3H).

To extend the Integrase Eraser platform, we engineered a new technology that we termed Swap, which uses integrase-based recombination to switch between sequence variants (isoforms) of the same gene. We built a prototype of Swap that enabled a switch from expression of wild type MED21 to expression of a mutant form of MED21 (mMED21) selectively in lateral roots (Fig. 4a). We employed a golden gate assembly approach to build our isoform target constructs, reminiscent of what we used to build our original constructs in Figure 1. The Swap target has a constitutive promoter (pUBQ10) flanked by integrase sites, positioned between the MED21 variants: wildtype MED21 marked with mTurquoise and a mutant form of MED21 marked with mScarlet. To avoid altering protein function through direct fusions, we instead introduced a P2A sequence between the MED21 and reporter sequences (Fig. 4b). The P2A peptide induces ribosomal skipping^28^, resulting in untagged versions of both MED21 variants and reporters.

**Figure 4.**
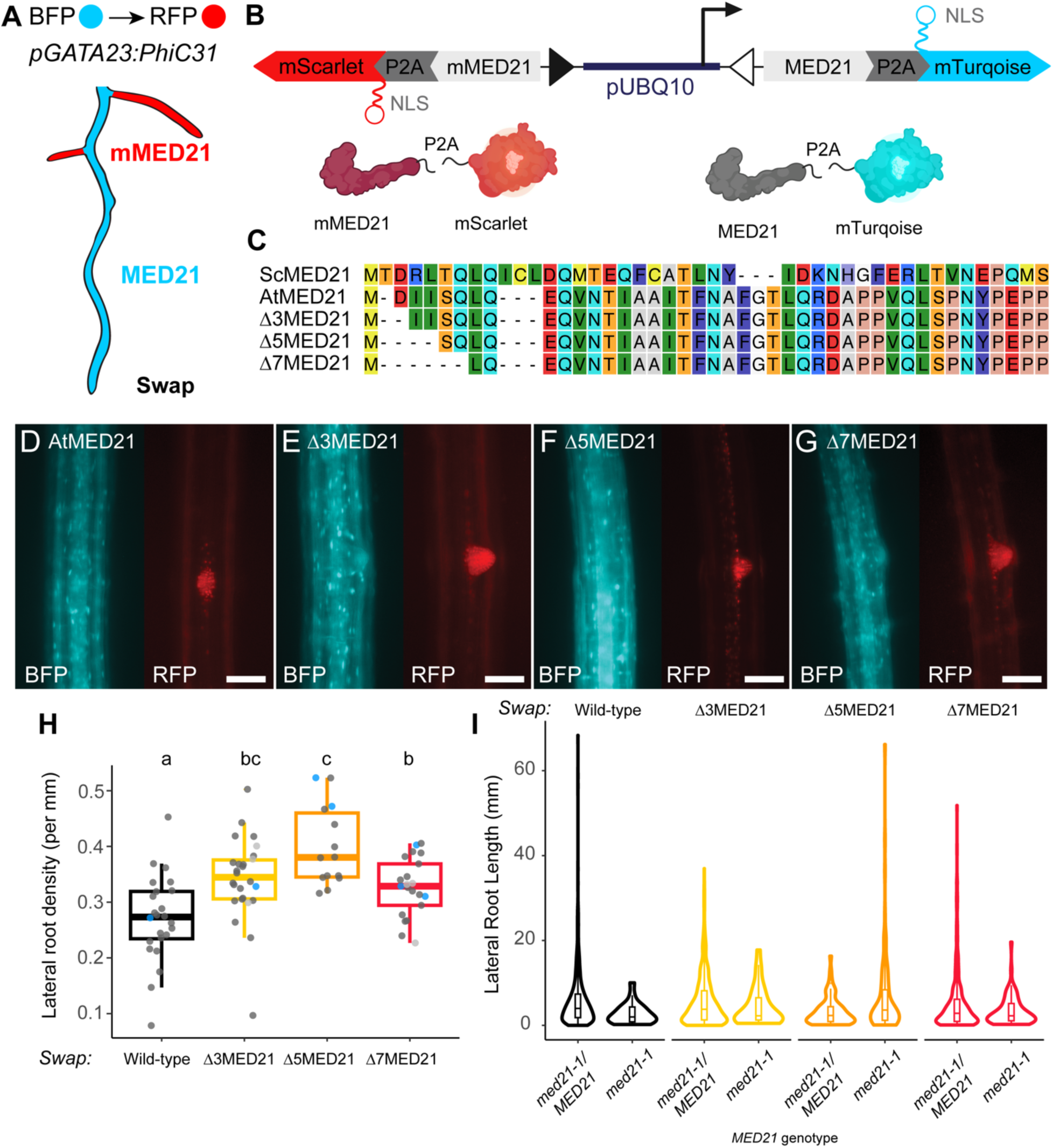
Engineering and rapid prototyping of the “hot-swap” isoform switch approach. **A.** Schematic of the predicted behavior for the MED21 Swap. Lateral root primordia will switch expression of the rescue construct from wild-type (blue) to mutant isoform (red). **B.** Design of the integrase target. The target is composed of two integrase sites (triangles) surrounding a constitutive promoter (pUBQ10), each MED21 isoform is fused to the P2A self-cleaving peptide sequence and a fluorescent reporter (mTurquoise or mScarlet). In the basal state, MED21 and mTurquoise are produced.

We selected mutant alleles of MED21 that are known to disrupt its binding with TPL (Figure 4C). Previous work in our lab has shown that deletions in the first seven amino acids of MED21 compromise TPL-binding activity and interfere with TPL-based repression^10^. Ectopic expression of any of these mutant *med21* isoforms in xylem pole pericycle cells competes with wild type MED21 resulting in increased lateral root density^10^. Here, we created four Swaps, in each wildtype MED21 is switched to a different *med21* mutant (Δ3MED21, Δ5MED21, Δ7MED21), in addition to the control which has a wild-type allele on either side of the switch (Fig 4C). We performed rapid prototyping in *Nicotiana benthamiana*, and found that switching of all Swap targets depended on the addition of the integrase (Figure S2A). Once exposed to integrase, Swap targets successfully switched, as indicated by expression of mScarlet.

We introduced the MED21 isoform switch targets into *med21-1/MED21, pGATA23::PhiC31* plants and performed fluorescence microscopy to identify primary transformant lines (T1) with specific switching to mScarlet in only the lateral roots (Figure 4E-G, Supplemental Figure 2B-E). We hypothesized that plants with mMED21 isoform switches should lead to increased lateral root density as was observed previously (Leydon et al., 2021). Additionally, the mMED21 isoforms should retain their activation activity in the Mediator complex leading to more normal lateral root lengths compared to the short lateral root phenotype observed in the MED21 Eraser experiments (Figure 2-3). Therefore, we quantified lateral root density in primary transformant mMED21 isoform switch lines (Lateral root density, Figure 4H). We observed that at mMED21 N-terminal mutation lines demonstrated higher lateral root density (Figure 4F) and that these lateral roots were similar in length to wild type (Figure 4I).

Once the PhiC31 integrase is expressed, it mediates inversion of the DNA between the integrase sites, inverting the promoter, and leading to production of mMED21 and mScarlet. **C.** Alignment of AtMED21 N-terminal mutants. **D-G.** Epifluorescence microscopy analysis of wild-type and mMED21 swap lateral root initiation in *Arabidopsis* primary transformants. Microscopy images were taken on a 20x objective, and the scale bar represents 50μm. **H.** Lateral root density (LRD per mm) and **I.** lengths of lateral roots (Lateral root length in mm) from primary transformant lines were quantified at 14 days post germination from the indicated Integrase Swap type. At least 15 independent primary transformant lines were tested for each swap type. Lines were transplanted to soil and PCR genotyped for *med21* genotype. In **H**, the LRD data points from med21 homozygous plants are colored in blue, while heterozygotes are colored dark gray, and ungenotyped samples are in light gray. Letters indicate significant difference (ANOVA and Tukey HSD multiple comparison test; p<0.001). in **I**, data is presented as violin plots with nested boxplots to demonstrate that medians and interquartile ranges are comparable across conditions.

## Conclusion

Essential genes are estimated to comprise approximately 20% of the genome^29^, and further techniques to study, modify and engineer these genes are needed to better understand and model their functions. Integrases are a promising tool, especially when compared to artificial microRNAs^30^ or CRISPR/Cas9-based tissue specific knock out (TSKO)^31^, which suffer from low efficiency. In addition, integrase-driven recombination events are highly precise, and less likely to produce the heterogeneity and diversity of repair-based mutations that result from Cas-based cleavage. While the tyrosine integrases such as Cre recombinase have been used more extensively in *Arabidopsis* and other plants to perform lineage tracing^32,33^, implement logic circuits^34^ and study an essential gene HAP2/GCS1^35^, there are several advantages that make serine integrases attractive. One clear advantage is that serine integrases bind to two distinct DNA binding sites making their modification of DNA irreversible. This is contrary to tyrosine integrases, such as FLP and Cre, which act on two identical sites and are able to catalyze both a forward and reverse reaction^36^. Another limitation of tyrosine integrase application in plants has been the observation of low efficiency activity due to CHH context methylation of the tyrosine recombinase sites^37^.

The current major drawback to the Integrase Eraser approach is the requirement for a heterozygote mutation in the gene of interest; however, large numbers of mutants from gametophyte^38^ and embryo lethal^19^ studies are ripe for reanalysis with such techniques. A more minor concern is that the high efficiency of integrase activity can lead to “overswitching” where recombination occurs in cells expressing only a background level of promoter expression. This potential challenge can be addressed by screening multiple lines, as is standard protocol for characterizing transgenic lines, and by using the integrase tuning toolkit described previously^20^ to attach protein degrons to the integrase, or RNA destabilization tags (DSTs) to its transcript. As a new suite of publicly-available tools that directly address the obstacles of pleiotropy and essentiality, Integrase Erasers should enable researchers experimental access to whole new swathes of the genome that have stood just out of reach

## Methods

### Cloning

Our cloning strategy was based on Golden Gate assembly using appropriate spacers and BsaI-HFv2 (NEB) as the restriction enzyme. The *Arabidopsis* MED21:HA sequence was amplified by PCR with appropriate Golden Gate restriction sites and the construction of integrase targets was performed by Golden Gate reaction in the modified pGreenII-Hygr vector containing compatible Golden Gate sites defined in Guiziou et al., 2022^20^. Enzymes for Golden Gate assembly were purchased from New England Biolabs (NEB, Ipswich, MA, USA). PCR was performed using 2X Q5 PCR master mix (NEB) and GoTaq master mix for colony PCR (Promega, Madison, WI, USA). Primers were purchased from IDT (Louvain, Belgium).Sequences were verified with Sanger sequencing by Azenta Life Sciences (Seattle, USA). Chemically-competent cultures of the E. coli strain DH5alphaZ1 (laciq, PN25-tetR, SpR, deoR, supE44, Delta(lacZYA-argFV169), Phi80 lacZDeltaM15, hsdR17(rK −, mK +), recA1, endA1, gyrA96, thi-1, relA1) were transformed with plasmid constructs containing kanamycin resistance. Transformed E. coli was grown in LB media (LB broth, Miller) with kanamycin (Millipore Sigma, 50 µg/mL).

### Western Blot

Protein was collected into 1.5ml tubes with one steel bead per tube (MN Beads Type D – 3mm steel beads, Machery-Nagel), snap frozen in liquid nitrogen, and homogenized on max settings on a tissue homogenizer (MM400, Retsch). Homogenized tissue was resuspended in 2x sample buffer, boiled for 10 minutes and spun down before being run on handmade 10% acrylamide SDS-PAGE gels, and western blotted with anti-HA-HRP antibodies from Roche/Millipore Sigma (RRID:AB_390917, REF-12013819001, Clone 3F10), or anti-Actin antibody (Abcam, ab197345). In all experiments equivalent mass of protein samples was used for extraction for compared sample types, i.e. root and shoots.

### Plant Growth

Arabidopsis seedlings were sown in 0.5 X Linsmaier and Skoog nutrient medium (LS) (Caisson Laboratories) and 0.8% w/v agar, stratified at 4 °C for 2 days, and grown in constant light at 22 °C. Phyto agar (PlantMedia/bioWORLD) was used when imaging seedlings and Bacto agar (ThermoFisher) was otherwise. For *Arabidopsis thaliana* experiments T2 plant lines harboring T-DNAs for MED21 (*med21-1*, WiscDsLox461-464K13) were grown on media supplemented with 25 µg/mL Glufosinate-ammonium (Oakwood Chemical, SC).

### Construction and selection of transgenic Arabidopsis lines

*Agrobacterium tumefaciens* strain GV3101 was transformed by electroporation, and subsequently grown in LB media with rifampin (Millipore Sigma, 50 µg/mL), gentamicin (Millipore Sigma, 50 µg/mL), any antibiotics carried on the specific plasmid(s), most often kanamycin (Millipore Sigma, 50 µg/mL). The floral dip method72 was used to generate integrase target lines in Col-0, and then used to introduce each integrase construct into these established target lines. For T1 selection: 120 mg of T1 seeds (∼2000 seeds) were sterilized using 70% ethanol and 0.05% Triton-X-100 and then washed using 95% ethanol. Seeds were resuspended in 0.1% agarose and spread onto 0.5X LS Bacto selection plates, using 25 µg/mL of kanamycin for target lines and 25 µg/mL kanamycin and 25 µg/mL hygromycin for lines with both the integrase and the target. The plates were stratified at 4 °C for 48 h then light pulsed for 6 h and covered for 48 h. They were then grown for 4–5 days. To select transformants, tall seedlings with long roots and a vibrant green color were picked from the selection plate with sterilized tweezers and transferred to a new 0.5X LS Phyto agar plate for characterization.

### Imaging of reporter and integrase lines

T1 seedlings for each line were grown 4–5 days after transformant selection. Each selected seedling was imaged at 20X magnification using an epifluorescence microscope (Leica Biosystems, model: DMI 3000) using the RFP (exposure 700 ms, gain 2) and CFP (exposure 700 ms, gain 2) channels. . Selected T1 seedlings were then transferred to soil, and at maturation T2 seeds were selected. For later generations, seedlings were sterilized similarly to T1s, stratified, plated on an LS agar plate, grown for 4–5 days, and characterized using the epifluorescence microscope as for T1. Confocal Imaging of the Eraser seedlings were performed using Nikon A1R HD25 laser scanning confocal microscope with 561 laser and 578-623 detector for RFP imaging. For the reporter lines, seedlings were scanned to find early-developed lateral roots. Imaging was processed using FIJI^39^. For each imaging, a Z-stack was recorded and a maximum average of the Z-stack in the RFP channel was generated.

### Root growth phenotyping

6-8 T2 seedlings from five selected *A. thaliana* lines were grown on antibiotic LS Phyto plates for 14 days. Each plate was scanned on days 10, 11, 12, 13, and 14 of growth using a flatbed scanner (Epson America, Long Beach, CA). Using FIJI, every lateral and primary root was measured in units of mm and every individual lateral root was counted. Then for each of the selected seedlings, we divided the number of lateral roots on the primary root by the root length. The data was graphed through use of R studio software. Fluorescence microscopy was used to identify and quantify the number of emerging lateral roots for every seedling.

For estradiol induction in T2s, antibiotic selection was performed as described in the method section about *A. thaliana* transgenic lines. Four days after transplanting Kanamycin, Hygromycin resistant seedlings onto 0.5X LS Phyto plates, the seedlings were imaged via microscopy in RFP channels with identical settings as described in the method section about integrase switch seedling characterization. Then the seedlings were transferred onto new 0.5X LS Phyto plates with 10 µM β-estradiol. Each seedling was imaged 24, 48, and 72 hours after transfer by both epifluorescence (for switching phenotype) and on a flatbed scanner as described above (for root growth phenotyping). A seedling was considered to exhibit a switch when any nuclear RFP was detected in the root tip region (manual scanning at least 10mm up each individual root tip). No distinction was made as to the proportion of cells switched, and unswitched seedlings were required to show no evidence of nuclear RFP.

### Protein alignments

The MED21 protein sequence was aligned to homologs using CLC Sequence Viewer 7 (QIAGEN, Aarhus, Denmark), a tree was constructed using a neighbor-joining method, and bootstrap analysis performed with 10,000 replicates.

### Quantification and statistical analysis

All quantification and statistical analyses were performed in R (4.4.2), and the corresponding code has been deposited into GitHub: https://github.com/achillobator/Hot-Swappable_Genetic_Switch

### Supporting information

**Supplemental Figures (S1-S2):**

**Supplemental Figure 1.** Characterization of independent MED21 iEraser lines, quantification of microscopy analysis.

**Supplemental Figure 2.** Characterization of swap targets in plants, in microscopy of rapid prototyping in *Nicotiana benthamiana*, and in *Arabidopsis thaliana* primary transformants.

**Supplemental Movie 1**. Time-lapse imaging of MED21 Eraser root growth phenotype.

## Acknowledgments

We thank current and former members of the Nemhauser group, including Dr. Sarah Guiziou, Janet Solano Sanchez, Benjamin Downing for constructive discussions. We thank Dr. Takato Imaizumi, Dr. Adam Steinbrenner, and Dr. Veronica Di Stillio for insightful suggestions. This work was supported by the NIH (R01-GM107084 and R35-GM148135-01 to JLN) and a Faculty Scholar Award from the Howard Hughes Medical Institute (to J.N.L.), A.R.L. was supported as a Simons Foundation Fellow of the Life Sciences Research Foundation.

## Author contributions

Conceptualization: ARL, JLN, Methodology: IJW, ARL, Software: IJW, ARL, Validation: IJW, ARL, Formal analysis: IJW, ARL, Investigation: IJW, CJM, ARL, Resources: JLN, Data Curation: IJW, ARL, Writing - Original Draft: IJW, Writing - Review & Editing, IJW, CJM, ARL, JLN, Visualization: IJW, CJM, ARL, Supervision: ARL, JLN, Project administration: ARL, JLN, Funding acquisition: ARL, JLN.

---

For Table of Contents Use Only

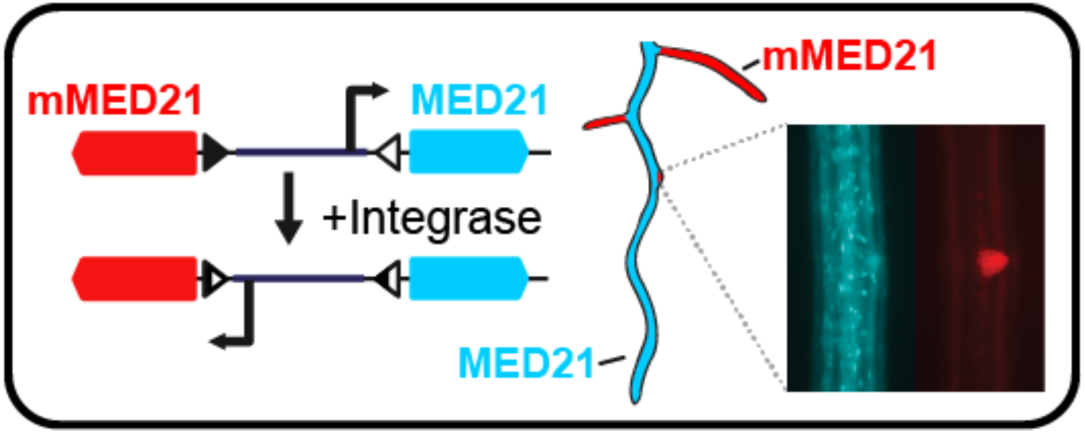

