## Supplemental Figures for "A Hot-Swappable Genetic Switch: Building an inducible and trackable functional assay for the essential gene MEDIATOR 21"

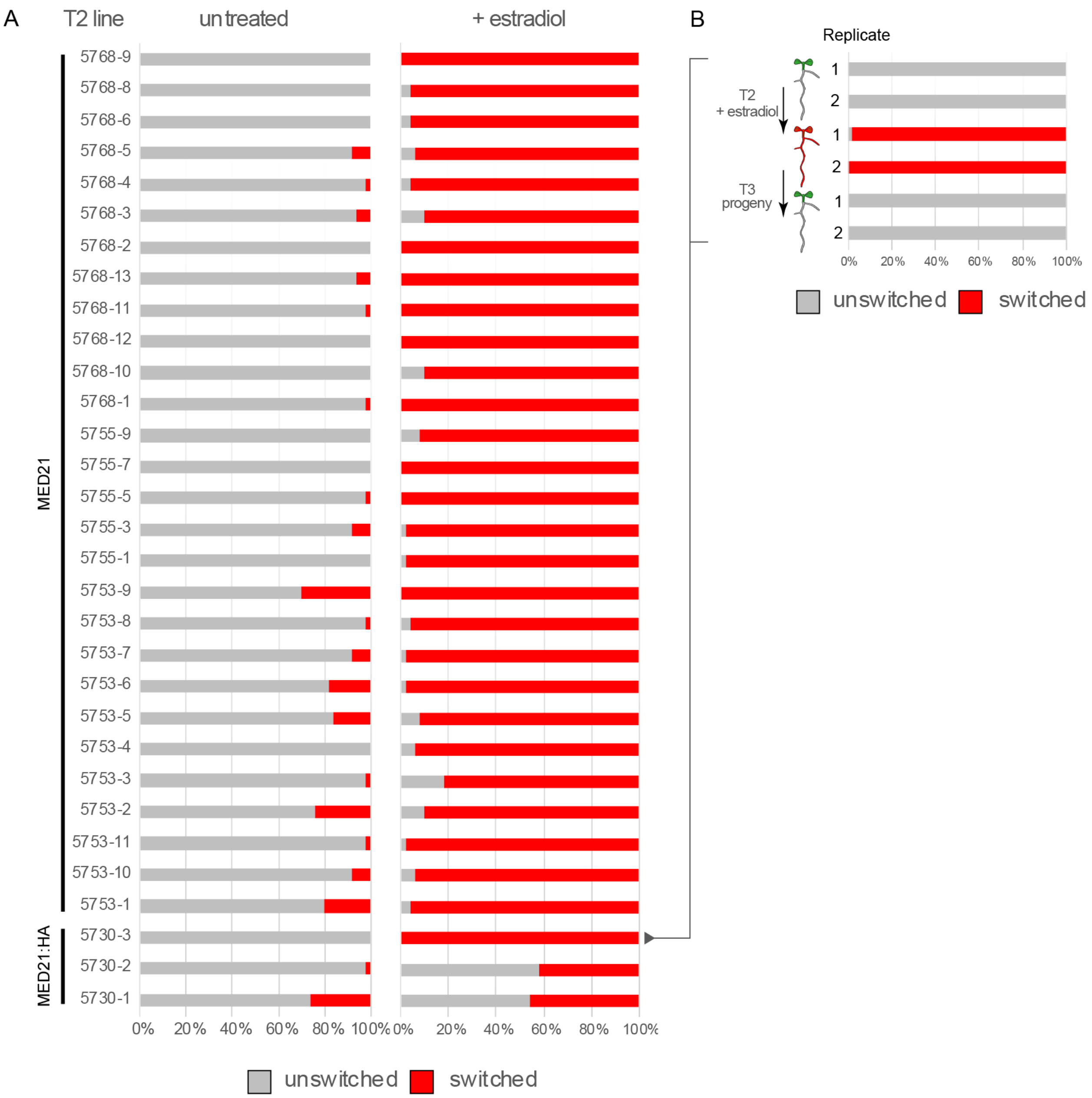


**S1. Characterization of independent MED21 iEraser lines. A.** Characterization of T2 seedlings. For the untagged MED21 eraser construct, we selected 28 T1 lines that were Kanamycin and Hygromycin resistant, and characterized 50 T2 seedlings per T1 line. Three MED21:HA eraser lines were included as controls (bottom three lines). Seedlings were seeded on Kanamycin and Hygromycin plates and grown for 5 days before being sprayed with 1$\mu$M estradiol. Plants were screened for RFP by fluorescence microscopy before and after application (3 days) and scored as either unswitched (gray) or switched (red). Data is presented as stacked bar graphs to highlight the proportion of the population undergoing a switch. B. Characterization of transmission of MED21:HA switch phenotypes to progeny. Switched seedlings for the highly specific line 5730-3 were transplanted (two replicates) to soil and allowed to produce seeds. These seeds were plated on media with no estradiol and screened for RFP by fluorescence microscopy. 50 seedlings were evaluated for each replicate of the experiment.

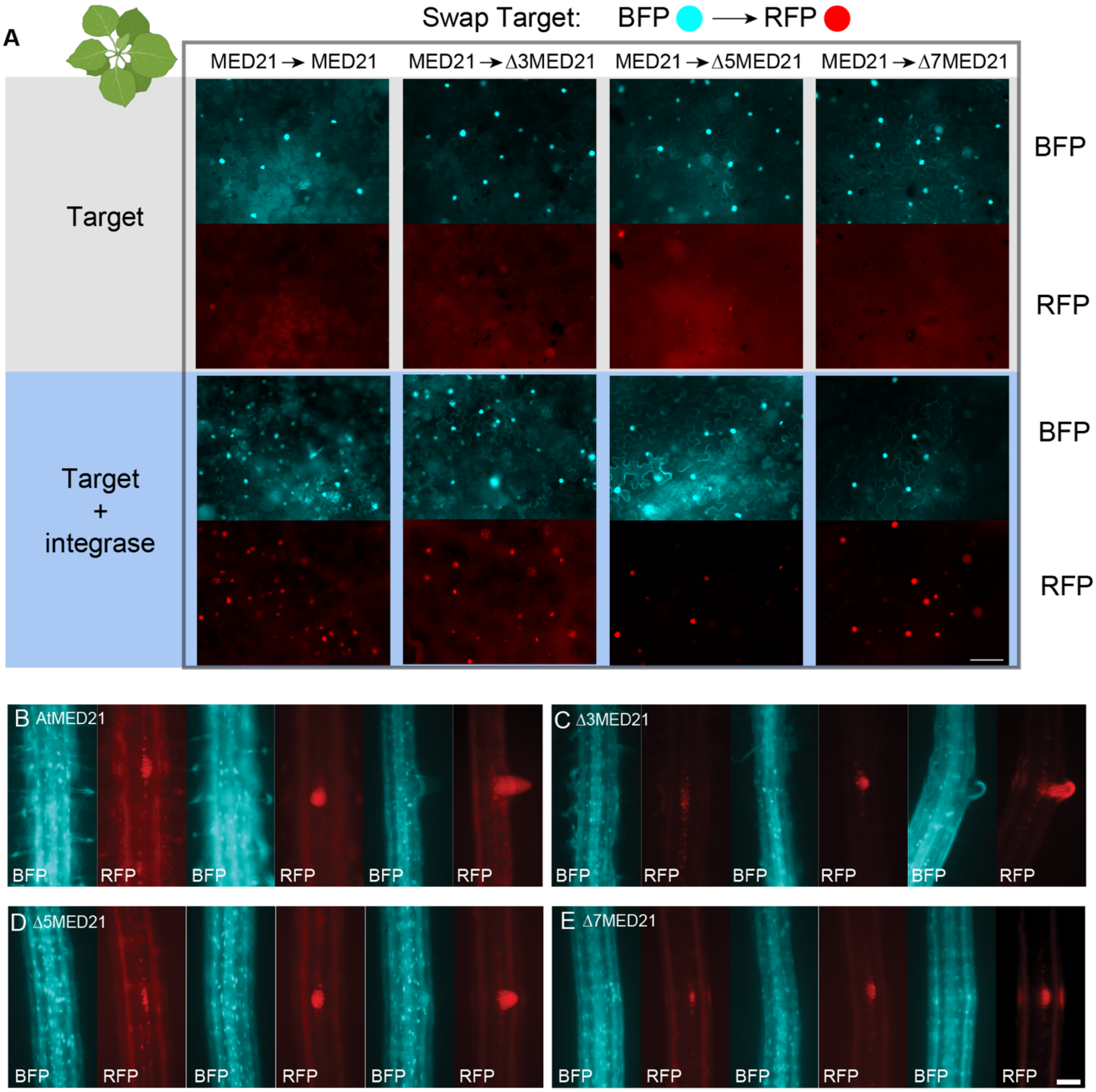


**S2. Characterization of swap targets in plants. A.** Rapid prototyping of the MED21 swap target in transient transfections of *Nicotiana benthamiana* at 2 days after injection. Top panel (gray backing) is the target alone. Bottom panel (blue backing) shows the impact of co-expressing the target with p35S:PhiC31. Microscopy images were taken on a 20x objective to allow a wide view of switching efficiency, the 50µm scale bar in the bottom right RFP channel applies to all images. **B-E.** Epifluorescence microscopy analysis of wild-type and MED21 swap lateral root initiation in *Arabidopsis* primary transformants. Microscopy images were taken on a 20x objective, and the 50µm scale bar in the bottom right RFP channel applies to all images.

**S3. Supplemental Movie S3. Time-lapse imaging of MED21 Eraser root growth phenotype.** Seedlings carrying the *pGATA23::PhiC31* driver were initially selected for Target and Driver constructs and T3 plants were confirmed to be homozygous for the *med21-1* mutation. Seedlings were grown for 5 days prior to hand transplantation on large square plates for time course analysis. All efforts were made to age and size match plants before transplantation. Images were taken every hour for 16 days, and the movie was cropped at the point at which the primary root neared the bottom of the plate. The genotypes are as follows: MED21 Eraser: 5750-5, 5754-9, MED21-HA Eraser: 5727-4, 5727-10, Control mTurquoise → mScarlet Target: 444.
